# TIR signaling promotes the interactions between EDS1/PAD4 and ADR1-L1 and oligomerization of ADR1-L1

**DOI:** 10.1101/2021.05.23.445317

**Authors:** Zhongshou Wu, Lei Tian, Xueru Liu, Yuelin Zhang, Xin Li

**Affiliations:** Michael Smith Laboratories, University of British Columbia, Vancouver, BC V6T 1Z4, Canada; Department of Botany, University of British Columbia, Vancouver, BC V6T 1Z4, Canada

## Abstract

Both plants and animals use nucleotide-binding leucine-rich repeat (NLR) immune receptors to perceive pathogens and trigger immunity. Toll/interleukin-1 receptor (TIR)-type plant NLRs (TNLs) require the lipase-like protein family members Enhanced Disease Susceptibility 1 (EDS1)/ Phytoalexin Deficient 4 (PAD4)/ Senescence-Associated Gene 101 (SAG101) and helper NLRs (hNLRs) for downstream signaling, the biochemical mechanisms of which remain unclear. Here, we report that TIR signaling promotes the association of EDS1 and PAD4 with hNLR ACTIVATED DISEASE RESISTANCE 1-Like 1 (ADR1-L1), and the oligomerization of ADR1-L1s for downstream immune activation and cell death.

## Introduction

Typical plant NLRs contain three functional domains. The C-terminal leucine-rich repeats (LRRs) are involved in effector recognition, self-repression and protein-protein interactions. The central nucleotide-binding (NB) region serves as an oligomerization platform and ATP/ADP-binding molecular switch (Jones et al., 2016). Based on the different N-termini, plant NLRs are grouped into three main subclasses: TNLs, coiled-coil (CC)-type NLRs (CNLs) and RESISTANCE TO POWDERY MILDEW 8 (RPW8)-like CC-type NLRs (RNLs) (Jubic et al., 2019). The TIR domains in TNLs have been shown to function as nicotinamide adenine dinucleotide (NAD^+^) hydrolase (NADases) (Wan et al., 2019; Horsefield et al., 2019). In *Arabidopsis thaliana*, there are two RNL sub-families, ACTIVATED DISEASE RESISTANCE 1 (ADR1) and N REQUIREMENT GENE 1 (NRG1). The ADR1 family contains ADR1, ADR1-Like 1 (ADR1-L1) and ADR1-Like 2 (ADR1-L2), with redundant functions. Similarly, the NRG1 family consists of the full-length NRG1A and NRG1B protein and an N-terminally truncated NRG1C. The full-length ADR1s and NRG1s function in parallel downstream of TNLs and in basal immunity with unequal redundancy (Dong et al., 2016; Wu et al., 2019; Castel et al., 2019; Jubic et al., 2019; Saile et al., 2020). Additionally, three lipase-like proteins are also required for TNL-mediated immunity, including EDS1, PAD4, and SAG101 (Wiermer et al., 2005). EDS1 interacts with PAD4 or SAG101 to form distinctive EDS1-PAD4 or EDS1-SAG101 heterodimers (Wagner et al., 2013), which work together with the ADR1s or NRG1s genetically, forming the EDS1-PAD4-ADR1s and EDS1-SAG101-NRG1s signaling modules downstream of TNLs (Lapin et al., 2019; Wu et al., 2019; Feehan et al., 2020). Besides TIR signaling, these two modules also contribute to plasma membrane localized receptor mediated immunity and basal defense (Tian et al., 2020; Pruitt et al., 2020). Recently, Sun et al. reported that NRG1A/1B interact with EDS1-SAG101 dimers in an effector-dependent manner to transduce TIR signals (Sun et al., 2020). However, how TIR signals are transduced by EDS1/PAD4/ADR1s was unclear.

## Results

During our previous analysis of *A. thaliana* hNLRs, we used *snc1* (*suppressor of npr1-1, constitutive 1*) and *chs3-2D* (*chilling sensitive 3, 2D*), two autoimmune TNL mutants, to establish the parallel relationships between the downstream EDS1-SAG101-NRG1 and EDS1-PAD4-ADR1 modules (Wu et al., 2019, Wu et al., 2021). To examine such relationship in the absence of constitutive defense activation, we generated a series of higher order mutants with CRISPR/Cas9, combining loss-of-function mutations from either different or the same modules in *A. thaliana* wild-type (WT) Col-0 background. All the newly generated mutants are indistinguishable from WT in morphology. We then challenged these plants with the avirulent bacterial pathogen *Pseudomonas syringae* pv. *tomato* (*P.s.t.*) DC3000 expressing either HopA1 or AvrRps4 effectors, which are recognized by TNL RPS6 (RESISTANT TO P. SYRINGAE 6) (Kim et al., 2009) or RPS4 (RESISTANT TO P. SYRINGAE 4) (Gassmann et al., 1999), and the virulent *Pseudomonas syringae* pv. *maculicola* (*P.s.m.*) ES4326. As shown in Figure S1A-C, mutants from the EDS1-PAD4-ADR1 modules, including *pad4-1* and *adr1 triple*, were similarly more susceptible compared to Col-0, and combining *adr1 triple* and *pad4-c1* mutations together failed to enhance their susceptibility further, supporting the functions of ADR1s and PAD4 in the same genetic module. However, mutants combining mutations in components from the two different modules, including *pad4-1 sag101-1*, *pad4-1 nrg1 triple*, *sag101-c1 adr1 triple*, and *adr1 nrg1 sextuplet*, are all more susceptible, supporting similar pathogen growth as *eds1-2* (Figure S1A-C). These results corroborate that EDS1-SAG101-NRG1 and EDS1-PAD4-ADR1 function as two distinct modules downstream of TNLs.

Genetically, ADR1s function together with EDS1/PAD4 in a signaling module. However, we were not able to reproducibly detect interactions between ADR1s and EDS1/PAD4 in co-immunoprecipitation (IP) experiments. Thus, we tested whether ADR1s associate with the EDS1-PAD4 heterodimer with a split luciferase complementation (SLC) assay. Among the three members of the ADR1 family, only ADR1-L1 expressed well and did not trigger HR in *Nicotiana (N.) benthamiana* (Figure S2). We therefore used ADR1-L1 in the follow-up experiments. As shown in Figure 1A-B, ADR1-L1 interacts with EDS1 (Figure 1A) as well as PAD4 (Figure 1B) by SLC (Figure S3). To confirm these interactions, we adopted the newly developed TurboID-based proximity labeling method, which allows TurboID-fused protein to biotinylate proximal and interacting proteins in the presence of biotin *in vivo* (Zhang et al., 2019), enabling detection of weak and transient protein-protein associations (Wu et al., 2021). As shown in Figure 1C, co-expression of EDS1-ZZ-TEV-FLAG and PAD4-ZZ-TEV-FLAG with ADR1-L1-HA-TurboID resulted in only a small amount of biotinylated EDS1-ZZ-TEV-FLAG and PAD4-ZZ-TEV-FLAG (Figure 1C). It is possible that the interactions between ADR1s and EDS1 or PAD4 require signals from upstream TIR/TNLs and the low levels of biotinylation of EDS1 and PAD4 by ADR1-L1-HA-TurboID are due to basal TNL activities in *N. benthamiana*. We therefore tested whether activation of TIR signaling can stimulate these interactions using the *A. thaliana* TIR-only protein RBA1 (RESPONSE TO HOPBA1), which triggers EDS1/PAD4-dependent immune responses (Nishimura et al., 2017). Interestingly, RBA1 treatment greatly increased the biotinylation of EDS1-ZZ-TEV-FLAG and PAD4-ZZ-TEV-FLAG by ADR1-L1-HA-TurboID (Figure 1C-D). However, the NADase-dead RBA1_E86A_ failed to induce a similarly enhanced biotinylation as WT RBA1 (Figure S4), suggesting that the NADase activity from TIR signaling enhances the interactions between ADR1-L1 and the EDS1-PAD4 heterodimer. In contrast, when EDS1-ZZ-TEV-FLAG and SAG101-3FLAG were expressed with ADR1-L1-HA-TurboID, only EDS1-ZZ-TEV-FLAG, but not SAG101-3FLAG, was biotinylated by ADR1-L1-HA-TurboID (Figure S5), supporting a specificity with EDS1-PAD4 dimer. Together, these data provide biochemical evidence to further support the distinct EDS1/PAD4/ADR1s and EDS1/SAG101/NRG1s modules.

**Figure 1.**
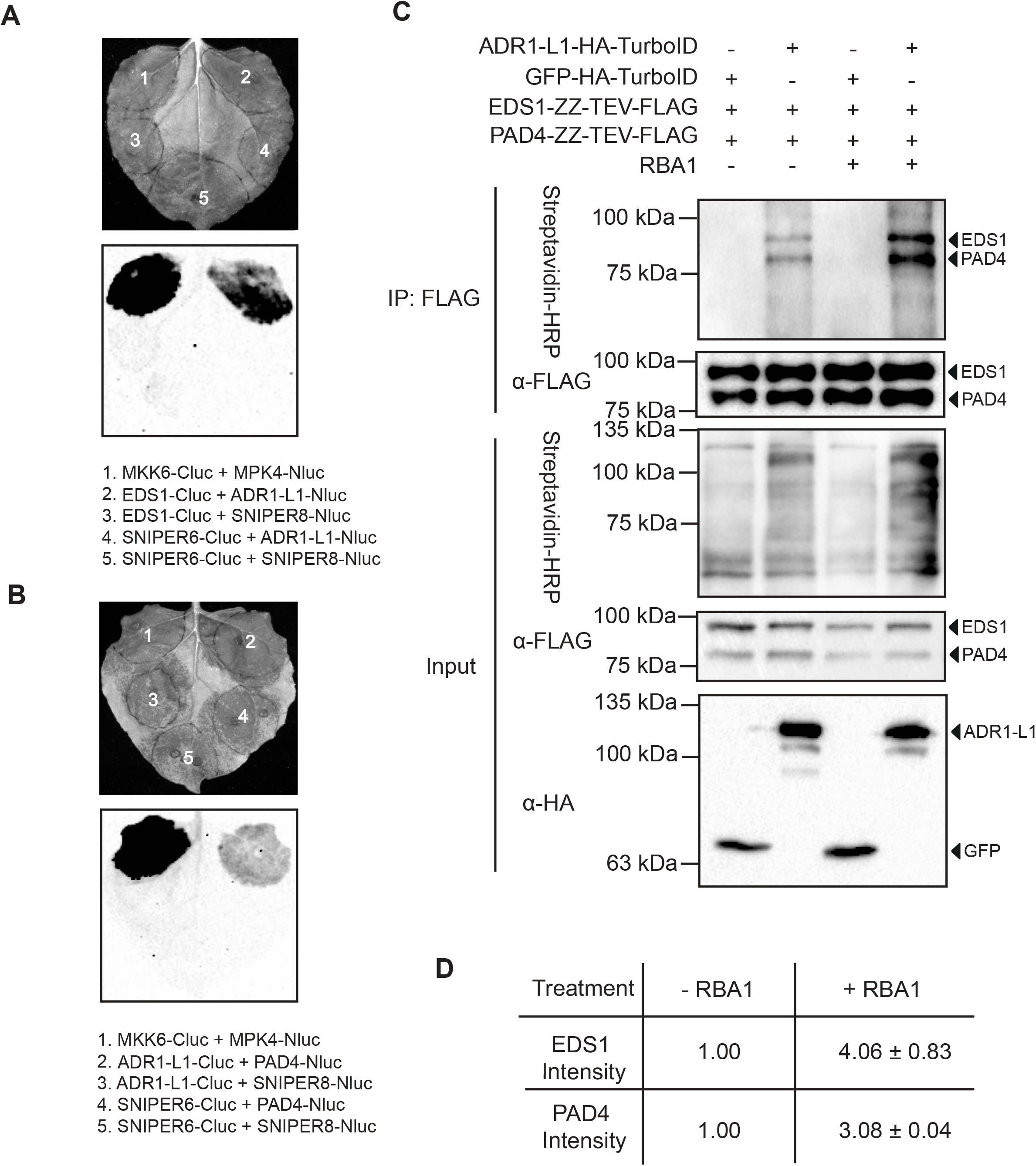
Activation of TIR signaling promotes the interaction between ADR1-L1 and the EDS1-PAD4 dimer. A-B. Interaction of ADR1-L1 with EDS1 (A) or PAD4 (B) as tested by split-luciferase complementation assay in *N. benthamiana*. The experiment was repeated three times with similar results. MPK4-Nluc and MKK6-Cluc were used as positive controls. The unpublished SNIPER6 and SNIPER8 are two immune-regulating E3 ligases isolated from *snc1*-influencing plant E3 ligase reverse (SNIPER) genetic screen, which were used as negative controls. C. Immunoprecipitation and biotinylation of EDS1-ZZ-TEV-FLAG and PAD4-ZZ-TEV-FLAG by ADR1-L1-HA-TurboID in *N. benthamiana* without or with RBA1 pre-treatment. Immunoprecipitation was carried out with anti-FLAG beads. The ZZ-TEV-FLAG-tagged proteins were detected using an anti-FLAG antibody. The HA-TurboID-tagged proteins were detected using an anti-HA antibody. The biotinylated proteins were detected using Streptavidin-HRP. Molecular mass marker in kiloDaltons is indicated on the left. The experiment was repeated three times with similar results. D. Quantification of EDS1-ZZ-TEV-FLAG and PAD4-ZZ-TEV-FLAG band intensity of (C) in Streptavidin-HRP blot. The numbers represent the normalized ratio between the intensity of the IP-enriched biotinylated protein band and the corresponding IP-enriched protein band in FLAG blot ± SD (n=3). Band intensity without RBA1 treatment was set to 1.

As oligomerization of NLRs is required for defense activation in both animal and plant systems (Bentham et al., 2017) and *N. benthamiana* NRG1 self-associates (Qi et al., 2018), we tested whether ADR1-L1 can interact with itself. As shown in Figure 2A, *in planta* association among ADR1-L1 proteins was observed in SLC. Such interaction was further confirmed with an IP experiment. When *N. benthamiana* leaves were co-infiltrated with the agrobacteria harbouring ADR1-L1-3HA and ADR1-L1-3FLAG, ADR1-L1-3FLAG was able to pull down ADR1-L1-3HA (Figure 2B), indicating that ADR1-L1 self-associates. Likewise, RBA1 induction considerably increased the amount of ADR1-L1-3HA being pulled down (Figure 2B-C). In contrast, RBA1_E86A_ was not able to increase ADR1-L1 self-association (Figure S6), suggesting that the upstream TIR signaling is responsible for enhancing the self-association of ADR1-L1. Previously it was shown that the CNL ZAR1 (HOPZ-ACTIVATED RESISTANCE 1) assembles into a pentameric complex (Wang et al., 2019), it will be interesting to determine whether ADR1-L1 also assembles into a pentamer upon activation.

**Figure 2.**
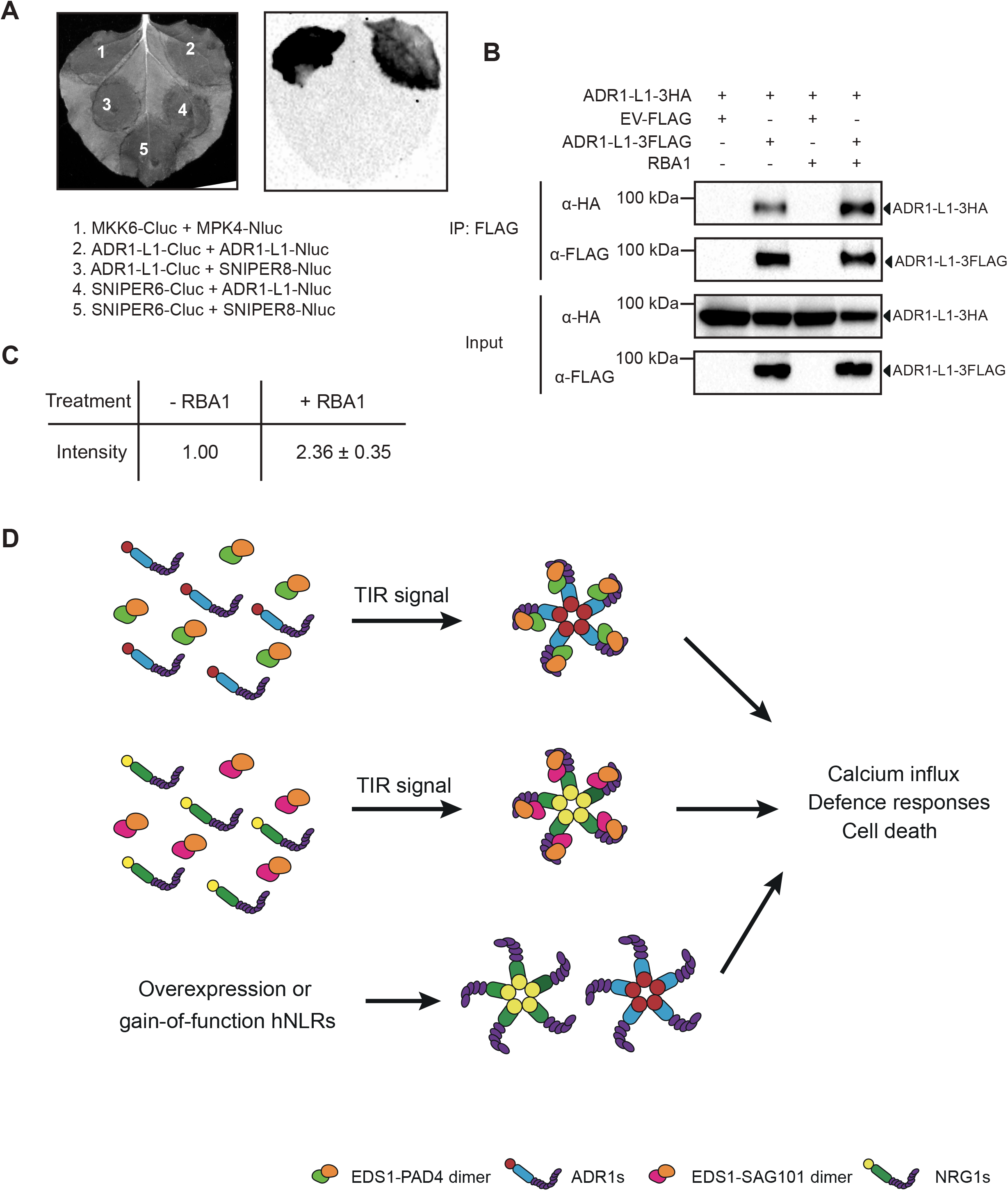
TIR signaling enhances the self-association of ADR1-L1. A. Self-association of ADR1-L1 as tested by split-luciferase complementation assay in *N. benthamiana*. The experiment was repeated three times with similar results. MPK4-Nluc and MKK6-Cluc were used as positive controls. SNIPER6 and SNIPER8 serve as negative controls. B. Immunoprecipitation of ADR1-L1-3HA by ADR1-L1-3FLAG in *N. benthamiana* without or with RBA1 pre-treatment. Immunoprecipitation was carried out with anti-FLAG beads. The 3FLAG-tagged proteins were detected using an anti-FLAG antibody. The HA-tagged proteins were detected using an anti-HA antibody. Molecular mass marker in kiloDaltons is indicated on the left. The experiment was repeated three times with similar results. C. Quantification of ADR1-L1-3HA band intensity of (B) in the anti-HA blot. The numbers represent the normalized ratio between the intensity of the ADR1-L1-3HA protein band by FLAG pull-down and the IP-enriched ADR1-L1-3FLAG protein band in FLAG blot ± SD (n=3). Band intensity without RBA1 treatment was set to 1. D. Working model of two defense modules upon activation of TIR signaling. TIR signaling leads to the generation of a product that can be perceived by either the EDS1-PAD4 or the EDS1-SAG101 dimers, which then triggers the assembly of EDS1-SAG101-NRG1 (Sun et al., 2020) and EDS1-PAD4-ADR1 complexes (current study) respectively to activate defense responses. The formation of the ADR1 or NRG1 pentameric resistosome complexes may serve as Ca^2+^ influx channels, resulting in downstream immune activation and cell death (Jacob et al., 2021). Overexpression or auto-active gain-of-function versions of the hNLRs can trigger self-oligomerizations and Ca^2+^ channel formation without the requirement of the lipase-like proteins.

The N-terminally truncated NRG1C can associate with the EDS1-SAG101 dimer and interfere with the EDS1-SAG101-NRG1 module function (Wu et al., 2021), implying that the part of NRG1 interacting with EDS1-SAG101 is likely through the C-terminal LRR and part of the NB domain. We therefore examined the EDS1-PAD4-ADR1 interactions with an N-terminally truncated ADR1. A construct overexpressing the truncated ADR1 similar to that of NRG1C (hereafter named as ADR1(1C)) was generated based on the protein sequence alignment (Figure S7 and Figure S8A) and was introduced into the *snc1* background, which contains a gain-of-function mutation in the *TNL SNC1* that results in ADR1s-dependent autoimmunity (Dong et al., 2016). As shown in Figure S8B-C, overexpression of *ADR1(1C)* partially suppressed *snc1*-mediated dwarfism and resistance to the oomycete pathogen *Hyaloperonospora arabidopsidis* (*H.a.*) Noco2. However, the transcript levels of *ADR1-L1* (Figure S8D) and *ADR1-L2* (Figure S8E) were not affected by the overexpression of *ADR1(1C)* (Figure S8F), excluding the possibility of suppression through gene silencing. The N-terminally truncated ADR1 likely acts as a dominant-negative form to interfere with the EDS1-PAD4-ADR1 module. In addition, when the HA-TurboID-ADR1(1C) was co-expressed with EDS1-ZZ-TEV-FLAG and PAD4-3FLAG in *N. benthamiana*, biotinylated EDS1-ZZ-TEV-FLAG and PAD4-3FLAG were detected (Figure S9A), suggesting that ADR1(1C) is in close proximity with EDS1 and PAD4. In agreement, addition of HA-TurboID-ADR1(1C) greatly reduced the SLC signals observed when EDS1-Cluc and ADR1-L1-Nluc were co-expressed (Figure S9B-C). Together, these data support that ADR1s likely associate with the EDS1-PAD4 dimer through their C-terminal NB-LRR region, although we cannot exclude the alternative explanation that the truncated ADR1 is in close proximity with EDS1/PAD4 via an interaction with endogenous full-length ADR1, rather than directly. Future structural and functional analyses are needed to resolve the detailed protein-protein interaction interfaces in the EDS1/PAD4/ADR1s module.

## Discussion

In summary, our study showed that activation of TIR signaling stimulates the interactions between ADR1-L1 and EDS1/PAD4 and self-association of ADR1-L1. Combining this with findings from two other studies which show that TIR signaling promotes the interaction between NRG1s and EDS1/SAG101 (Qi et al., 2018; Sun et al., 2020), a conceptual model for the roles of EDS1/PAD4/SAG101 and the hNLRs in TIR signaling is proposed (Figure 2D). Upon recognition of pathogen effectors, TIR/TNL receptors are activated, leading to the generation of an NADase product(s), which is subsequently recognized by EDS1-PAD4 and EDS1-SAG101. Recognition of this signal molecule(s) is proposed to stimulate the interaction of EDS1-PAD4 and EDS1-SAG101 with the hNLRs, leading to self-association of hNLRs and formation of hNLR oligomeric complexes. In contrast, overexpression or auto-active versions of hNLRs are capable of self-association (Saile et al., 2020), leading to EDS1-independent defence activation (Qi et al., 2018; Wu et al., 2019) (Figure S10, Figure 2D). According to two very recent reports (Bi et al., 2021; Jacob et al., 2021), similar to ZAR1, the hNLR oligomers can also form Ca^2+^ channels on the plasma membrane to activate downstream immune responses and cell death. Therefore, the hNLRs serve as receptors for the ligand-bound EDS1-PAD4 and EDS1-SAG101 heterodimers for TIR signaling. Upon activation, they may form ZAR1 resistosome-like pentameric rings that can serve as Ca^2+^ influx channels to turn on immune responses and cell death. Future investigations on the nature of the signaling molecule produced by the NADase activity of the TIR domain and how it interacts with the lipase-like proteins are needed for full understanding of the TIR signaling pathway in plants.

## Methods

### Plant growth conditions

*A. thaliana* and *N. benthamiana* plants were grown at 22 °C under a long day condition of 16 hr light/8 hr dark regime.

### Construction of plasmids

Overexpression and HA-TurboID tagged *ADR1*(*1C*) constructs were made in *pBASTA* and *pBASTA-N-HA-TurboID* vectors, respectively. *EDS1*, *PAD4* and *ADR1-L1* were cloned into *pCambia 1300-Cluc* or *-Nluc*. *PAD4* was cloned into *pBASTA-3FLAG*. *ADR1-L1* was cloned into *pBASTA-HA-TurboID*, *pCambia1300-3FLAG* and *pCambia1300-3HA*. All primers used are listed in Supplemental Table 1. *pCambia1305 EDS1-ZZ-TEV-FLAG*, *pCambia1305 PAD4-ZZ-TEV-FLAG*, *pBASTA GFP-HA-TurboID* were described in previous studies (Wu et al., 2021). The CRISPR/Cas9 constructs for knocking out *PAD4* and *SAG101* were described in previous studies (Wu et al., 2021).

### *A. thaliana* transformation

The binary constructs were introduced into *Agrobacterium tumefaciens* GV3101 by electroporation and subsequently transformed into *A. thaliana snc1* background by the floral-dipping method (Clough & Bent, 1998). Transformants were selected on soil by spraying BASTA (Glufosinate ammonium). The co-segregation analysis was then performed with the transformants in T_2_ generations to make sure the suppression phenotypes were due to the transgene overexpression.

### Pathogen infection assay

Pathogen infection assays were carried out as described previously (Li et al., 2001). In brief, two-week-old soil-grown seedlings were sprayed with 10^5^ *H.a.* Noco2 conidia spores per ml water. The plants were transferred to a humid chamber at 18 °C of 12 hr light/12 hr dark. After 7 days, sporulation was quantified using a hemocytometer. For bacterial infections, four-week-old plants were infiltrated with bacterial solution at designated concentrations using a blunt-end syringe. Leaf discs were collected and ground at 0- and 3-day post infiltration. Colony-forming units (CFU) were calculated after incubation on LB plates with appropriate antibiotic selection.

### TurboID-based proximity labeling in *N. benthamiana*

TurboID-based proximity labeling assay was performed as described previously (Wu *et al.*, 2020; Zhang *et al.*, 2019). In brief, *N. benthamiana* leaves were infiltrated with agrobacteria containing *HA-TurboID*, *3FLAG* and *ZZ-TEV-FLAG* tagged constructs. At 24 hpi, agrobacteria expressing RBA1 constructs as well as the biotin were infiltrated. A similar amount of leaf samples was harvested at 36 dpi, followed by immunoprecipitation and western blot analysis.

### Protein extraction, co-immunoprecipitation, and western blot analysis

For protein level examination, 0.1 g *N. benthamiana* leaves were collected and extracted by extraction buffer (100 mM Tris-HCl pH 8.0, 0.2% SDS and 2% β-mercaptoethanol). Loading buffer was added to each protein sample and boiled for 5 min before loading.

Co-immunoprecipitation assay was performed as previously described (Wu et al., 2020). In brief, about 2.0 g *N. benthamiana* leaves expressing the indicated proteins were harvested at 36 hpi and ground into fine powder with liquid nitrogen. Extraction buffer contains 25 mM Tris-HCl pH 7.5, 300 mM NaCl, 5 mM MgCl_2_, 0.15% Nonidet P-40, 10% Glycerol, 1 mM PMSF, 1× Protease Inhibitor Cocktail (Roche; Cat. #11873580001), and 10 mM DTT. For testing the interaction between ADR1-L1 with EDS1-PAD4 and the self-association of ADR1-L1, 20 μM dATP and 5 mM CaCl_2_ were added to the extraction buffer to be able to consistently observe the interactions, as NLRs are ATPases and the maintenance of their oligomeric form requires the presence of ATP/dATP (Wang et al., 2019). The FLAG-tagged proteins were immunoprecipitated using 20 μl M2 beads (Sigma Cat. #A2220) and biotinylation was detected by Streptavidin-HRP (Abcam Cat. # ab7403). The anti-HA antibody was from Roche (Cat. #11867423001). The anti-FLAG antibody was from Sigma (Cat. #F1804). Protein abundance was quantified using ImageJ (https://imagej.nih.gov/ij/).

### Split luciferase complementation assay

Agrobacteria expressing the luciferase C-terminus (C-luc) and N-terminus (N-luc)-fused proteins were co-infiltrated into four-week-old *N. benthamiana* leaves, and the infiltrated leaves were treated with 1 mM luciferin after 48 hpi. Each bacterial strain was diluted to a final concentration of OD_600_ = 0.2.

### Expression analysis

About 0.05 g of four-week-old soil-grown plant tissue was collected, and RNA was extracted using an RNA isolation kit (Bio Basic; Cat#BS82314). ProtoScript II reverse transcriptase (NEB; Cat#B0368) was used to generate cDNA. Real-time PCR was performed using a SYBR premix kit (TaKaRa, Cat#RR82LR). The primers used are listed in Supplemental Table 1.

### Statistical analysis

Statistical analysis was carried out with one-way ANOVA followed by Tukey’s post hoc test. The Scheffé multiple comparison was applied for testing correction. Normality test for all data was done in SPSS. Statistical significance was indicated with different letters. *p* values and sample numbers (n) were claimed in figure legends.

## Author contributions

ZW and LT, Data curation, Validation, Investigation, Methodology, Writing - original draft, Project administration; X Liu, Validation, Methodology; YZ and X Li, Conceptualization, Data curation, Formal analysis, Supervision, Funding acquisition, Writing—original draft, Project administration, Writing - revisions. All authors reviewed the manuscript.

## Acknowledgments

Drs. Jonathan Jones, Pingtao Ding, Yu Ti Cheng and Shengyang He are cordially thanked for sharing of *Pseudomonas* strains. Dr. Marc T. Nishimura is thanked for generous sharing of the RBA1 constructs. Ms. Solveig van Wersch is sincerely thanked for careful reading of the manuscript. This work was financially supported by CFI-JELF, the Natural Sciences and Engineering Research Council of Canada (NSERC) Discovery Program, NSERC-CREATE PRoTECT program, and the WD Cooper Memorial Fund from UBC. L.T. is partly supported by a China Scholarship Council (CSC) scholarship.

---

**Figure S1.**
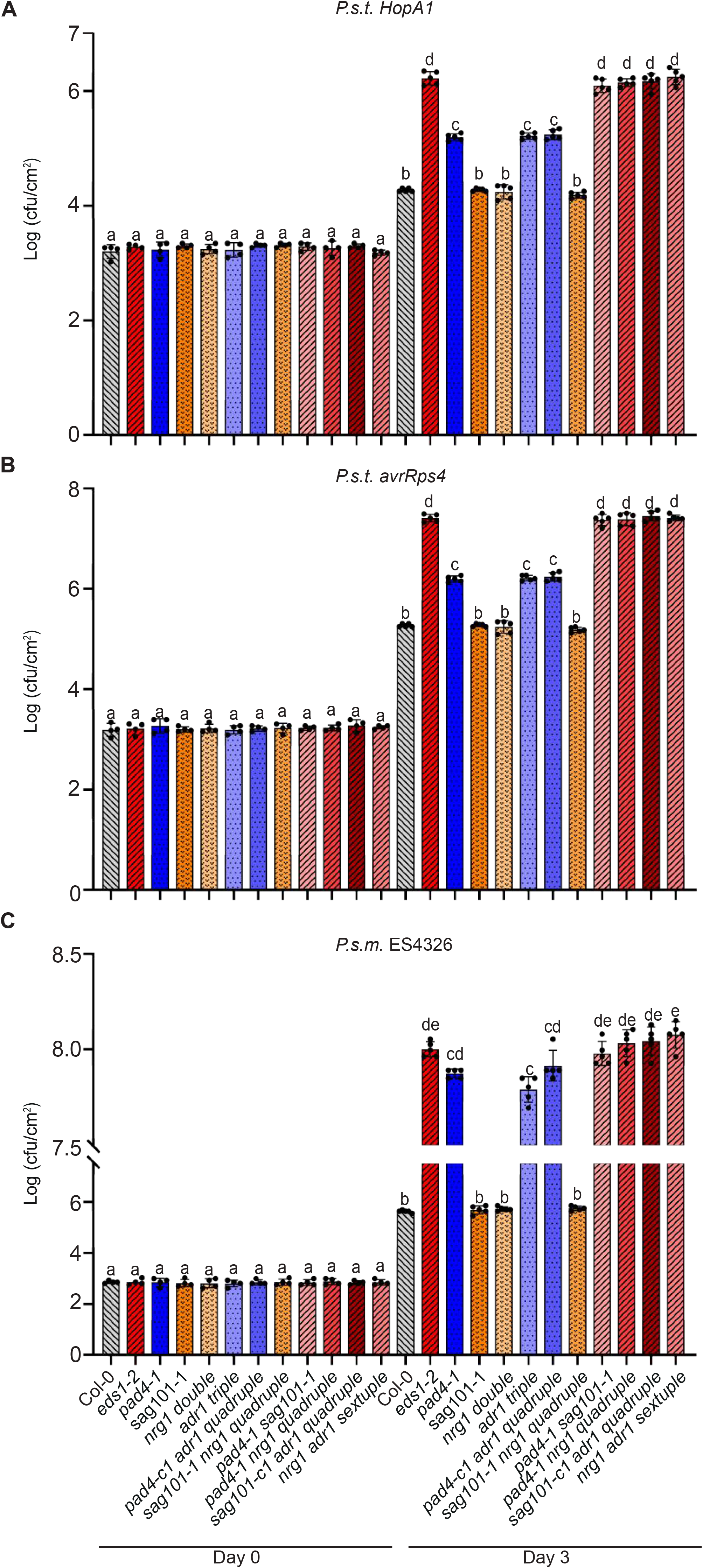
The EDS1-PAD4-ADR1 module functions in parallel with the EDS1-SAG101-NRG1 module. A-C. Growth of *P.s.t. HopA1* (A), *avrRps4* (B) and *P.s.m.* ES4326 (C) in four-week-old leaves of the indicated genotypes at 0 dpi and 3 dpi, with bacterial inoculum of OD_600_ = 0.0001. Statistical significance is indicated by different letters (*p* < 0.01). Error bars represent means ± SD (n=4 for day 0, n=5 for day 3). Three independent experiments were carried out with similar results.

**Figure S2.**
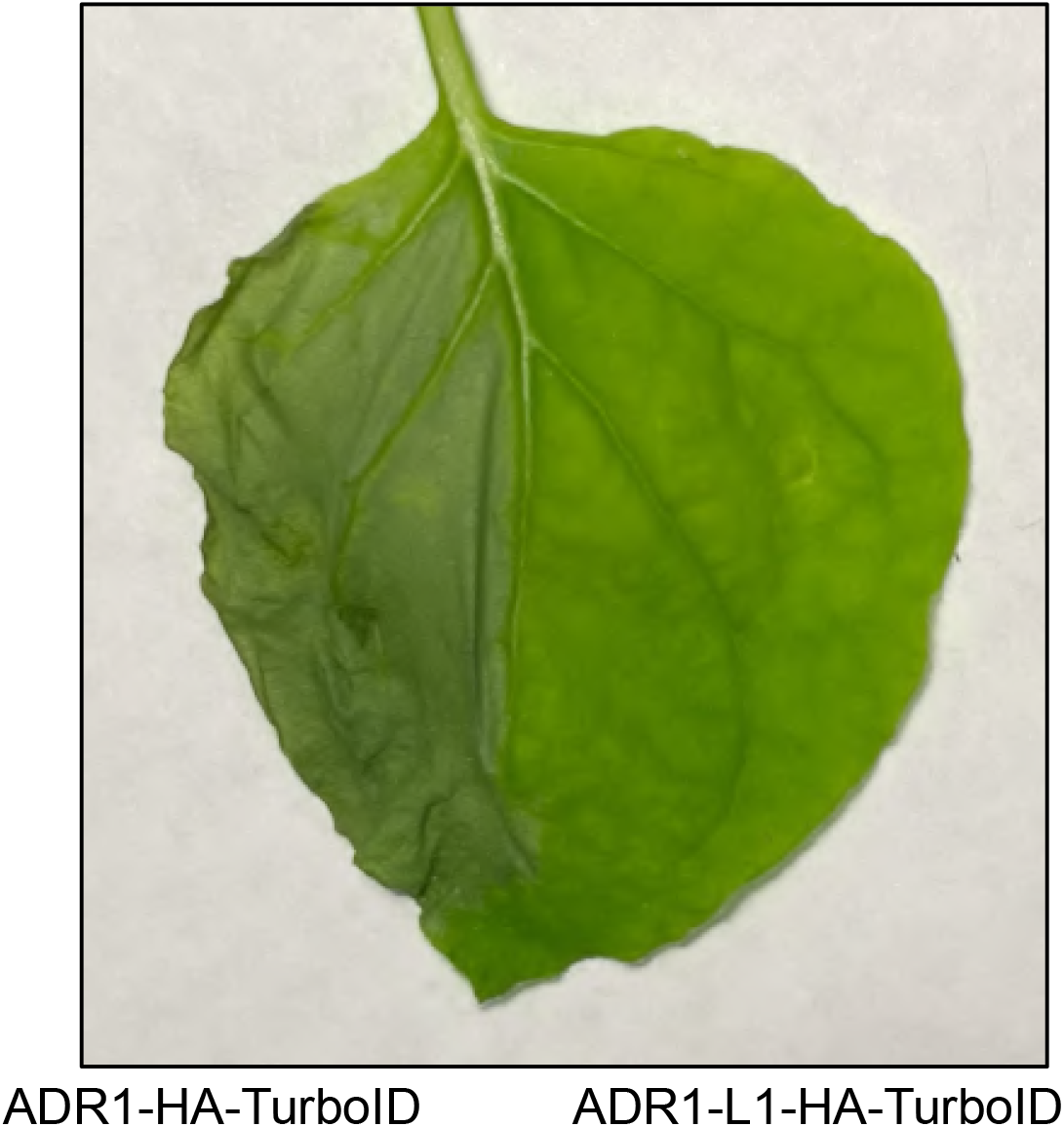
ADR1-HA-TurboID, not ADR1-L1-HA-TurboID, causes cell death in *N. benthamiana*. HR in the *N. benthamiana* leaves expressing ADR1-HA-TurboID (left) or ADR1-L1-HA-TurboID (right). Photos were taken at 36 hpi. Three independent experiments were carried out with similar results.

**Figure S3.**
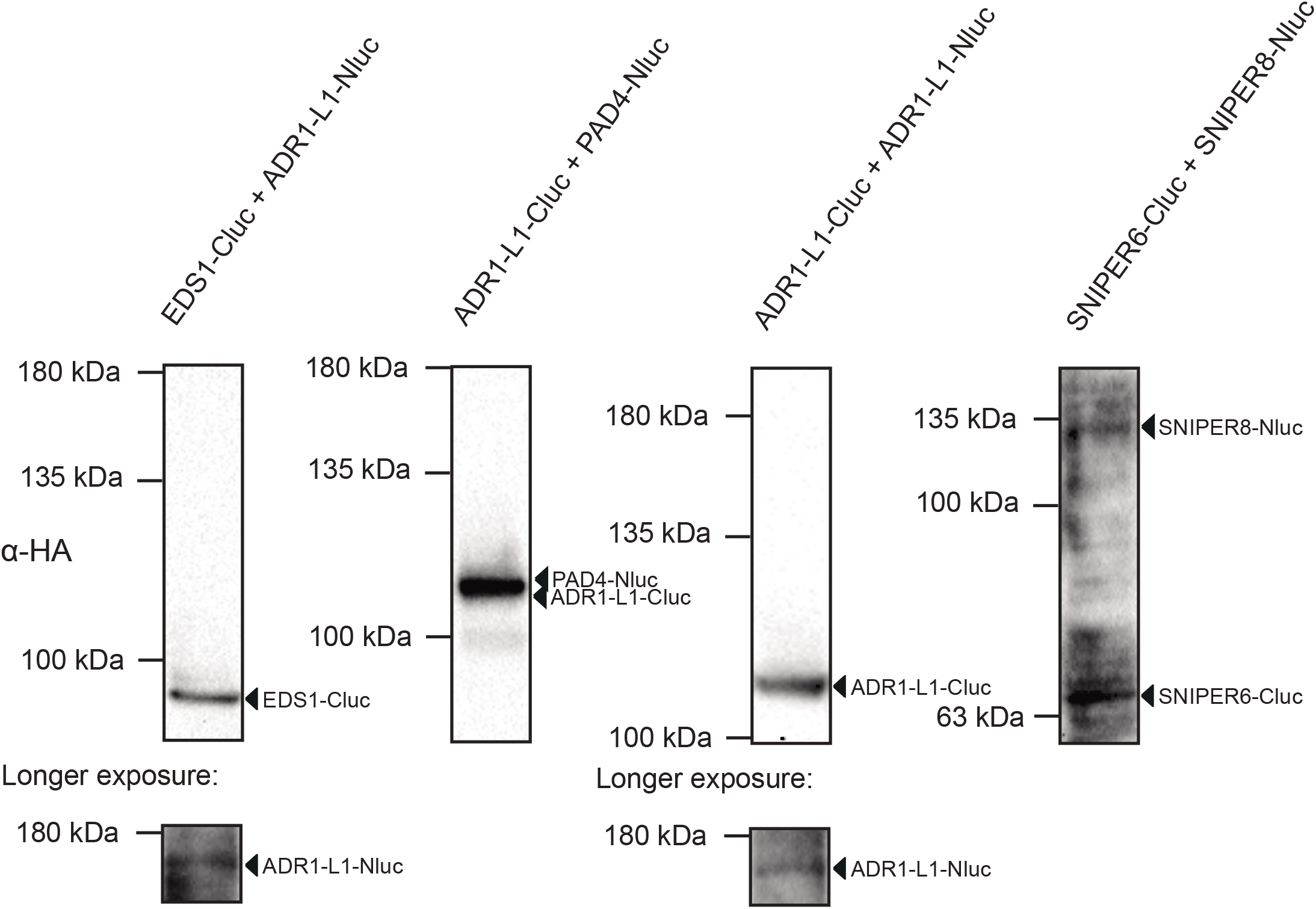
The protein expressions in SLC assays in *N. benthamiana*. Western blots showing the expression of EDS1-Cluc, ADR1-L1-Nluc, PAD4-Nluc, ADR1-L1-Cluc, SNIPER6-Cluc and SNIPER8-Nluc. Molecular mass marker in kiloDaltons is indicated on the left. The molecular weights of PAD4-Nluc (111.7 kDa) and ADR1-L1-Cluc (109 kDa) are too close to separate.

**Figure S4.**
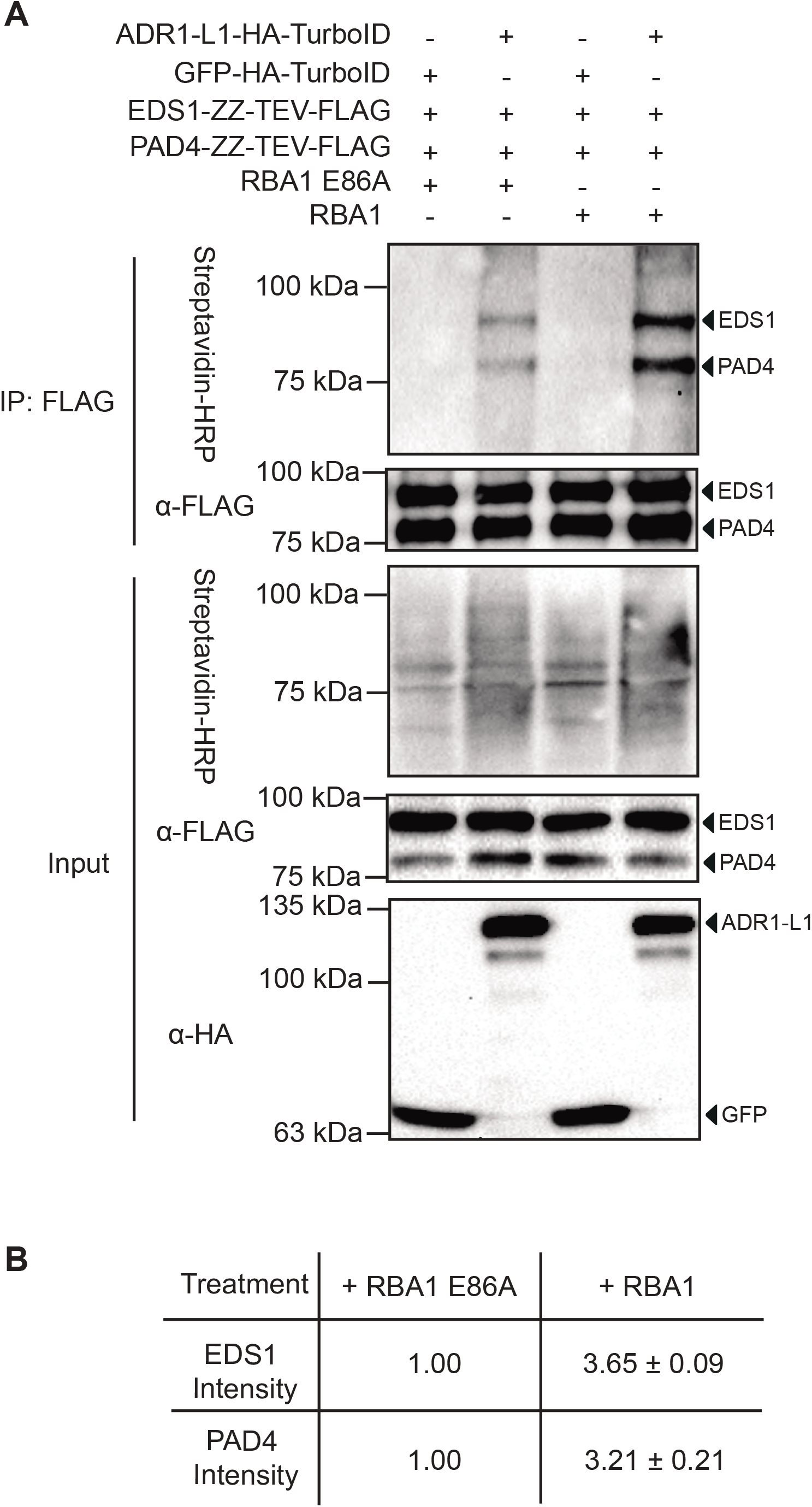
The NADase dead RBA1_E86A_ fails to enhance the association between ADR1-L1 with EDS1/PAD4. A. Immunoprecipitation and biotinylation of EDS1-ZZ-TEV-FLAG and PAD4-ZZ-TEV-FLAG by ADR1-L1-HA-TurboID in *N. benthamiana* with RBA1 or RBA1_E86A_ pre-treatment. Immunoprecipitation was carried out with anti-FLAG beads. The ZZ-TEV-FLAG-tagged proteins were detected using an anti-FLAG antibody. The HA-TurboID-tagged proteins were detected using an anti-HA antibody. The biotinylated proteins were detected using Streptavidin-HRP. Molecular mass marker in kiloDaltons is indicated on the left. The experiment was repeated twice with similar results. B. Quantification of EDS1-ZZ-TEV-FLAG and PAD4-ZZ-TEV-FLAG band intensity of (A) in Streptavidin-HRP blot. The numbers represent the normalized ratio between the intensity of the IP-enriched biotinylated protein band and the corresponding IP-enriched protein band in FLAG blot ± SD (n=2). Band intensity with RBA1_E86A_ treatment was set to 1.

**Figure S5.**
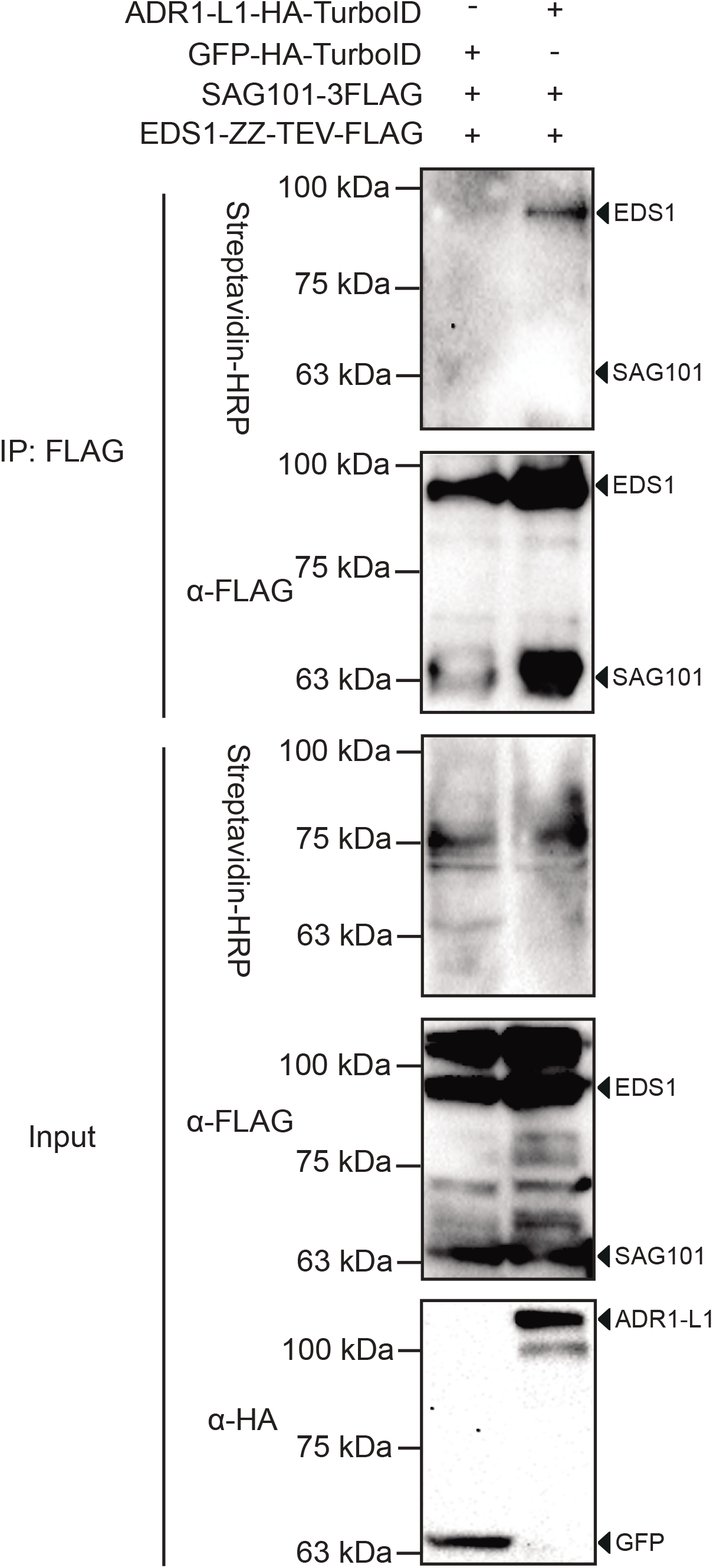
EDS1-ZZ-TEV-FLAG, but not SAG101-3FLAG, is biotinylated by ADR1-L1-HA-TurboID. A. Immunoprecipitation and biotinylation of EDS1-ZZ-TEV-FLAG and SAG101-3FLAG by ADR1-L1-HA-TurboID in *N. benthamiana*. Immunoprecipitation was carried out with anti-FLAG beads. The ZZ-TEV-FLAG and 3FLAG-tagged proteins were detected using an anti-FLAG antibody. The HA-TurboID-tagged proteins were detected using an anti-HA antibody. The biotinylated proteins were detected using Streptavidin-HRP. Molecular mass marker in kiloDaltons is indicated on the left. The experiment was repeated twice with similar results.

**Figure S6.**
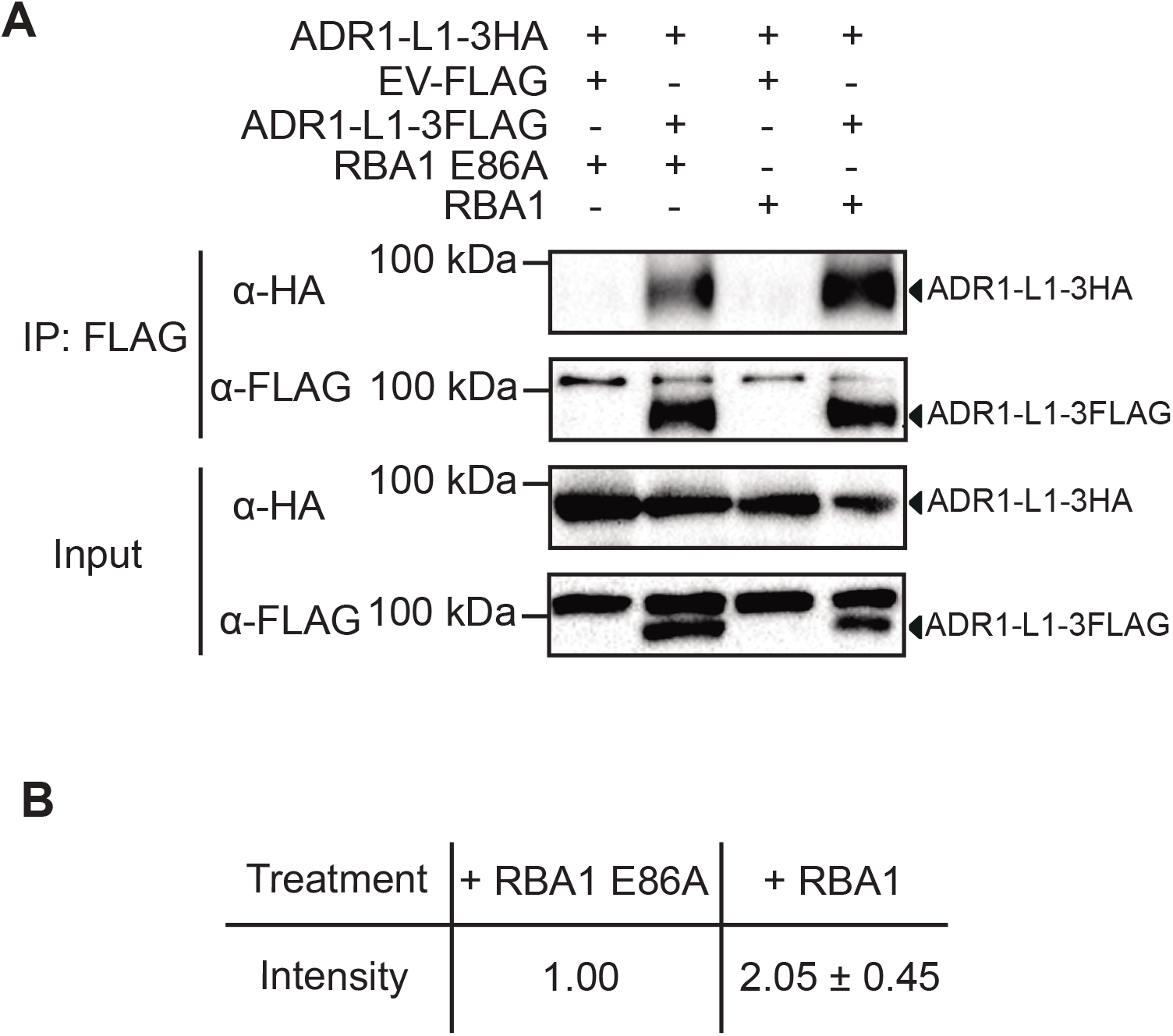
RBA1_E86A_ did not enhance ADR1-L1 self-association as with WT RBA1. A. Immunoprecipitation of ADR1-L1-3HA by ADR1-L1-3FLAG in *N. benthamiana* with RBA1 or RBA1_E86A_ pre-treatment. Immunoprecipitation was carried out with anti-FLAG beads. The 3FLAG-tagged proteins were detected using an anti-FLAG antibody. The HA-tagged proteins were detected using an anti-HA antibody. Molecular mass marker in kiloDaltons is indicated on the left. The experiment was repeated twice with similar results. B. Quantification of ADR1-L1-3HA band intensity of (A) in the anti-HA blot. The numbers represent the normalized ratio between the intensity of the ADR1-L1-3HA protein band by FLAG pull-down and the IP-enriched ADR1-L1-3FLAG protein band in FLAG blot ± SD (n=2). Band intensity without RBA1_E86A_ treatment was set to 1.

**Figure S7.**
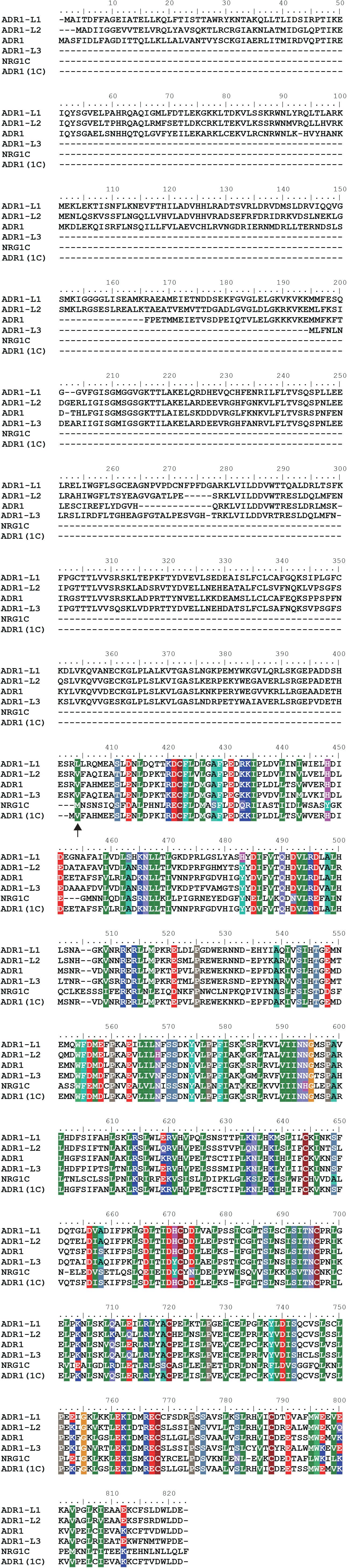
Protein alignment of *A. thaliana* ADR1-L1, ADR1-L2, ADR1, ADR1-L3, NRG1C and NRG1C-type truncated ADR1 (ADR1(1C)). Protein sequences were obtained from TAIR. Arrow indicates the truncation location on ADR1.

**Figure S8.**
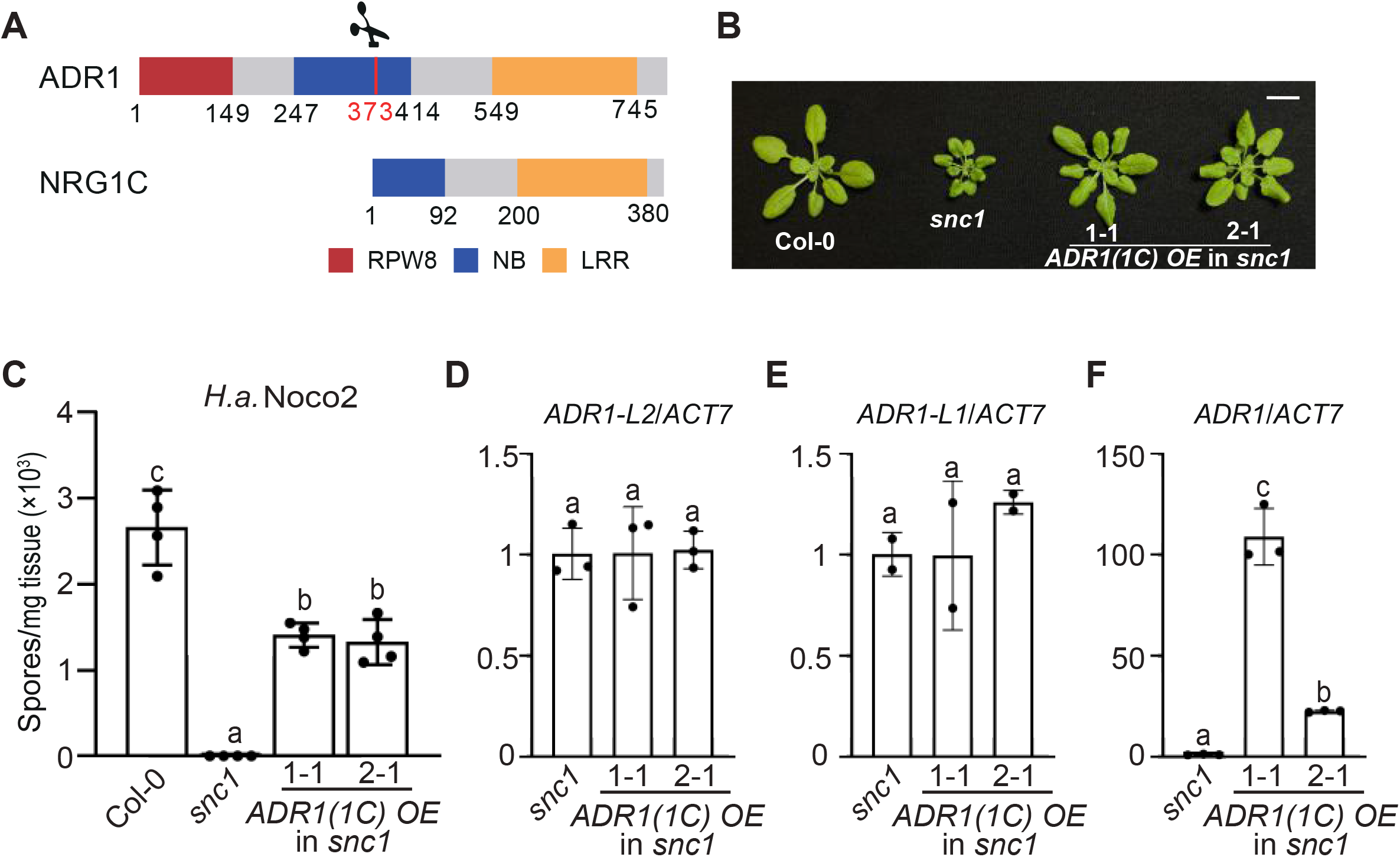
Overexpression of *ADR1*(*1C*) partially suppresses *snc1*-mediated dwarfism/autoimmunity. A. Protein domain diagram of NRG1C and ADR1. Numbers represent amino acid positions relative to the translation start sites. Scissor and red line indicate the truncation location on ADR1 for generating ADR1(1C). B. Morphology of four-week-old soil-grown plants of Col-0, *snc1* and two independent transgenic lines of *ADR1(1C) OE* into *snc1* background. C. Quantification of *H.a.* Noco2 sporulation in the indicated genotypes at 7 days post inoculation (dpi) with 10^5^ spores per ml water. Statistical significance is indicated by different letters (*p* < 0.01). Error bars represent means ± SE (n=4). Three independent experiments were carried out with similar results. D-F. Expression levels of *ADR1-L2* (D), *ADR1-L1* (E) or *ADR1* (F) in the indicated genotypes as determined by RT-PCR (*snc1* serves as control whose transcript level was set at 1.0). The *ADR1* transcripts measured include the ones for both the native full-length *ADR1* and the truncated *ADR1(1C)* as the primers used for RT-PCR can amplify both. Statistical significance is indicated by different letters (*p* < 0.01). Error bars represent means ± SE (n = 3 for D and F; n=2 for E). Two independent experiments were carried out with similar results.

**Figure S9.**
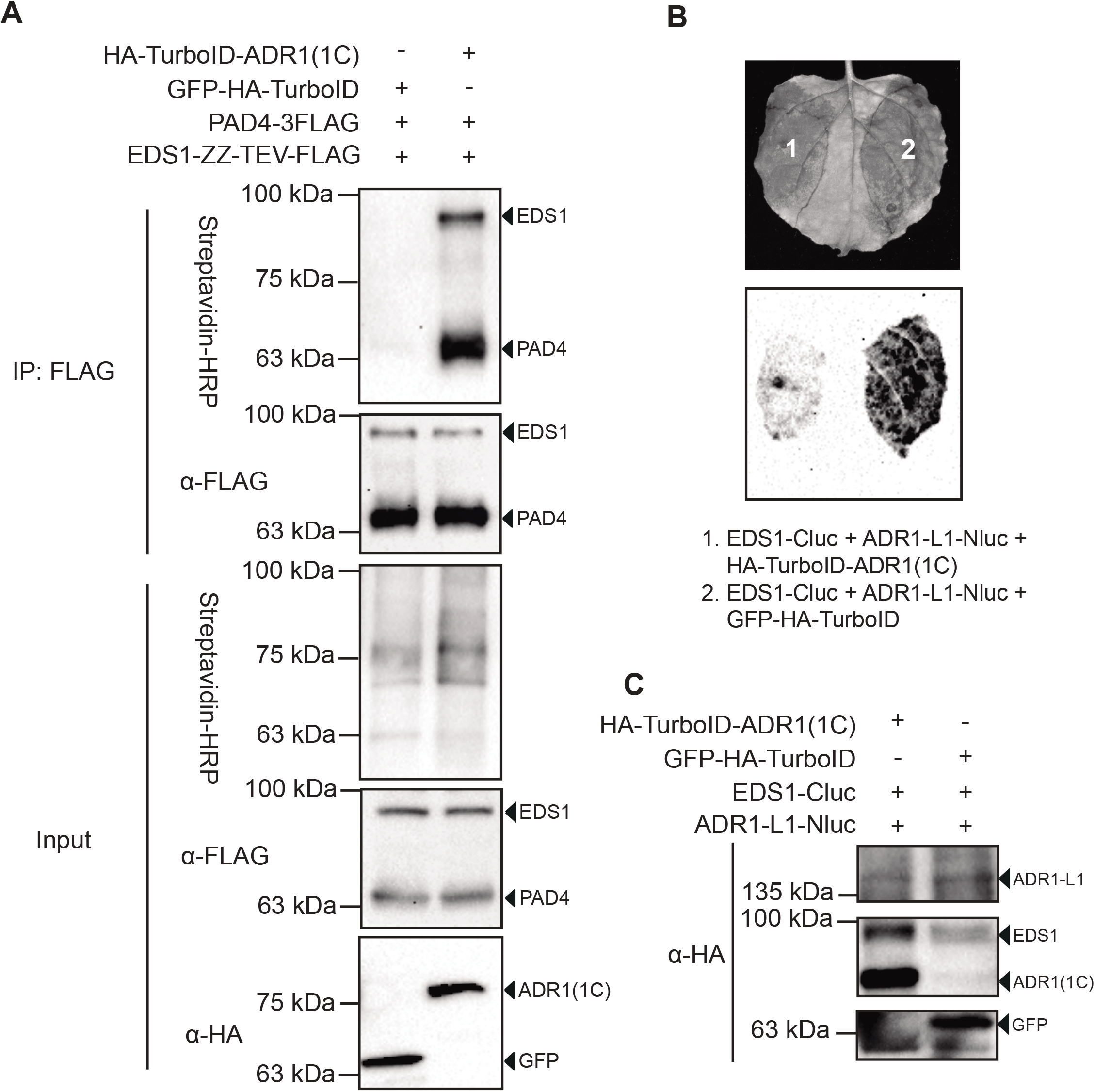
ADR1(1C) interacts with EDS1-PAD4 dimer. A. Immunoprecipitation and biotinylation of EDS1-ZZ-TEV-FLAG and PAD4-3FLAG by HA-TurboID-ADR1(1C) in *N. benthamiana*. Immunoprecipitation was carried out with anti-FLAG beads. The ZZ-TEV-FLAG and 3FLAG-tagged proteins were detected using an anti-FLAG antibody. The HA-TurboID-tagged proteins were detected using an anti-HA antibody. The biotinylated proteins were detected using Streptavidin-HRP. Molecular mass marker in kiloDaltons is indicated on the left. The experiment was repeated three times with similar results. B. Interaction of ADR1-L1 with EDS1 as tested by split-luciferase complementation assay in *N. benthamiana* in the presence of HA-TurboID-ADR1(1C) (Left) or GFP-HA-TurboID (Right). The experiment was repeated three times with similar results. C. Western blot showing the protein expression in (B). Molecular mass marker in kiloDaltons is indicated on the left.

**Figure S10.**
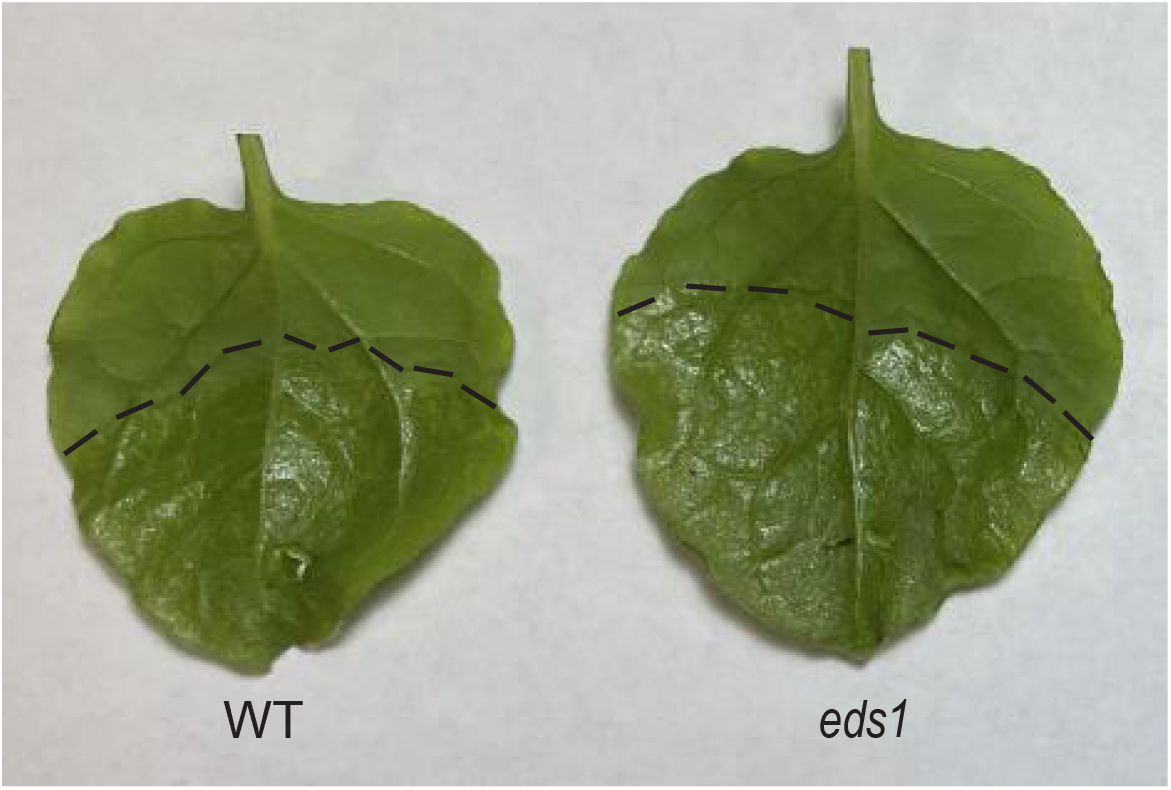
ADR1-3FLAG causes cell death in WT and *eds1 N. benthamiana*. HR in the *N. benthamiana* leaves expressing ADR1-3FLAG in WT (Left) or *eds1* (Right) *N. benthamiana*. Photos were taken at 36 hpi. Three independent experiments were carried out with similar results.

**Table S1.**
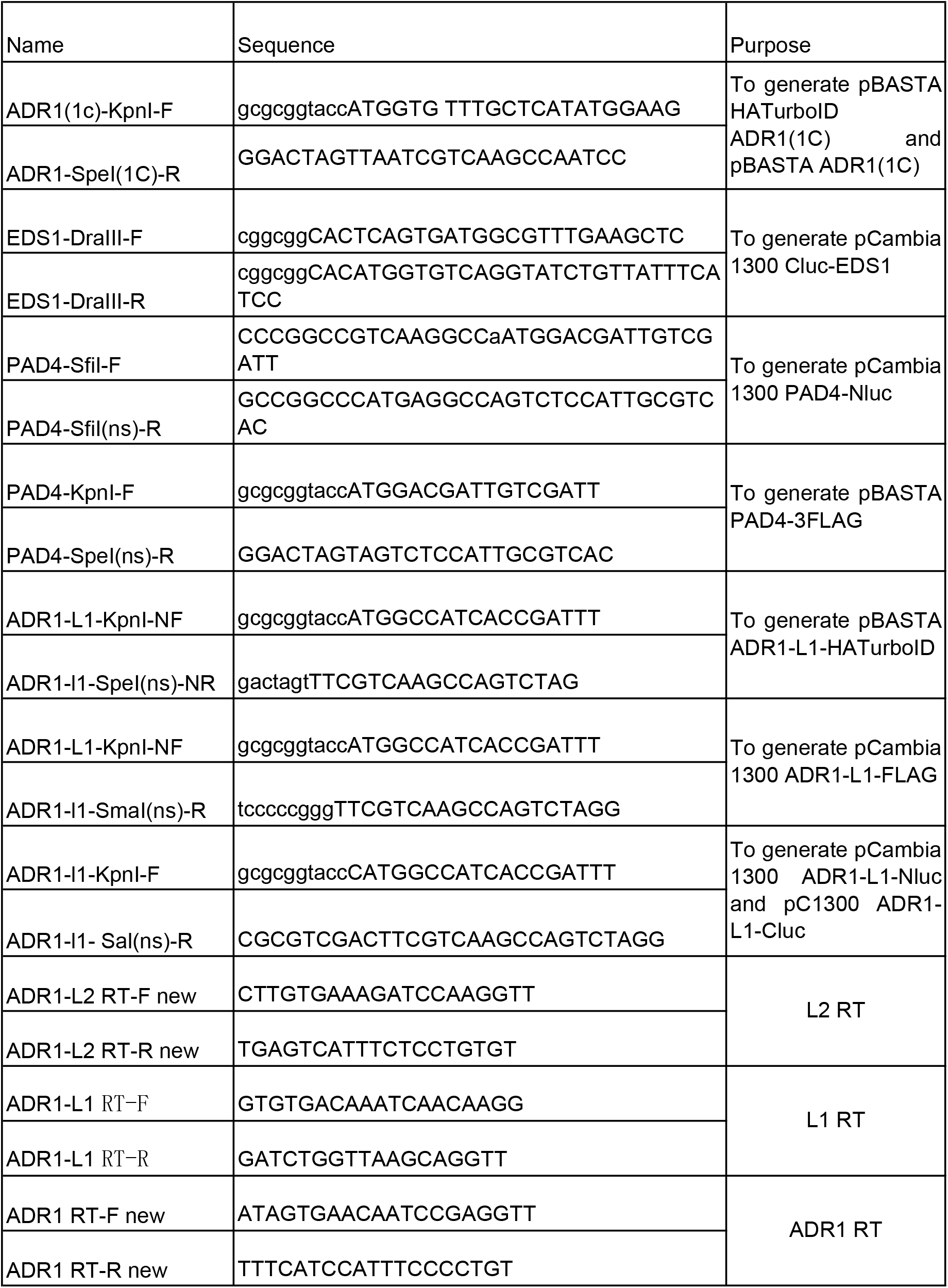
The list of primers used in this study.

